# Heme is crucial for medium-dependent metronidazole resistance in clinical isolates of *C. difficile*

**DOI:** 10.1101/2020.11.18.388959

**Authors:** Ilse M. Boekhoud, Igor Sidorov, Sam Nooij, Céline Harmanus, Ingrid M.J.G. Bos-Sanders, Virginie Viprey, Bill Spittal, Emma Clark, Kerrie Davies, Jane Freeman, Ed J. Kuijper, Wiep Klaas Smits, On behalf of the COMBACTE-CDI Consortium and the European Study Group for *Clostridioides difficile*

**Affiliations:** Department of Medical Microbiology, Leiden University Medical Center, Leiden, The Netherlands; Netherlands Centre for One Health, The Netherlands; Center for Microbiome Analyses and Therapeutics, Leiden University Medical Center, Leiden, The Netherlands; Healthcare Associated Infections Research Group, School of Medicine, University of Leeds, Leeds, United Kingdom; National Institute for Public Health and the Environment, Bilthoven, The Netherlands

## Abstract

Until recently, metronidazole was the first-line treatment for *Clostridioides difficile* infection and it is still commonly used. Though resistance has been reported due to the plasmid pCD-METRO, this does not explain all cases. Here, we investigate resistance to metronidazole in a collection of clinical isolates of *C. difficile*. We find that nearly all isolates demonstrate a heme-dependent increase in the minimal inhibitory concentration for metronidazole, which in some cases leads to isolates being qualified as resistant (MIC > 2 mg/L). Moreover, whole genome sequence analysis reveals a single nucleotide polymorphism in the heme responsive gene *hsmA*, which defines a metronidazole resistant lineage of PCR ribotype 010 / multilocus sequence type 15 isolates that also includes pCD-METRO containing strains. Together our data demonstrate that heme is crucial for medium-dependent metronidazole resistance in *C. difficile*.

## Introduction

*Clostridioides difficile* is a gram-positive, anaerobic enteropathogen capable of causing a *C. difficile* infection (CDI) upon disruption of the normal intestinal microbiota by for instance antimicrobial therapy. ^1, 2^ It is the primary cause of nosocomial diarrhea, but is also found in cases of community-acquired disease. ^2, 3^ Although the use of antibiotics is a risk factor for CDI, antimicrobials are also used to treat the infection. Until recently, metronidazole was considered the drug-of-choice for treatment of mild CDI. Though vancomycin and fidaxomicin are currently indicated as first-line therapeutics for the treatment of CDI, ^4, 5^ metronidazole is still commonly used. ^6, 7^

Our understanding of the mechanisms of resistance to metronidazole in *C. difficile* is still limited. For Clostridia, studies are complicated by reports of unstable, inducible metronidazole resistance which is lost upon removal of antibiotic pressure or after the strain undergoes freeze-thawing cycles. ^8, 9^ Recently, however, we have demonstrated that metronidazole resistance in diverse strains of *C. difficile* can be mediated by the plasmid pCD-METRO through a mechanism that is not yet understood. ^10^ Notably, the presence of pCD-METRO explains at least part of independently reported cases of metronidazole resistance. ^11-13^ However, strains that lack pCD-METRO can still be resistant to metronidazole. For instance, we have previously identified a pCD-METRO negative strain that demonstrated medium-dependent metronidazole resistance. ^10^ This suggests that other, potentially chromosomal, determinants contribute to resistance.

Information on pathways that could contribute to resistance comes from laboratory strains with evolved resistance to metronidazole. Using a laboratory-evolved PCR ribotype (RT) 027 strain exhibiting stable metronidazole resistance, mutations were identified in genes affecting electron transport and iron utilisation, but the individual contribution of these mutations to the resistance phenotype was not further investigated. ^14^ More recently, a mutator strain defective in DNA mismatch repair was evolved in the presence of metronidazole. ^15^ The study discovered mutations in a gene encoding an iron transporter (*feoB1*) in metronidazole resistant strains and showed that sequential mutations in *nifJ* (encoding the pyruvate-ferredoxin oxidoreductase, PFOR), *xdh* (encoding xanthine dehydrogenase) or *iscR* (encoding an iron-sulfur cluster regulator) could further increase metronidazole resistance. ^15^ Though studies with laboratory evolved strains are informative, it is unclear how these findings translate to metronidazole resistant strains isolated from subjects outside the laboratory.

Here, we leverage the potential of strains collected within a project to develop a detailed understanding of the epidemiology and clinical impact of CDI across Europe (COMBACTE-CDI) that were investigated for metronidazole resistance. Four strains demonstrated stable metronidazole resistance, but lacked pCD-METRO. We show that there is a heme-dependent increase in the minimal inhibitory concentration (MIC) for metronidazole across PCR ribotypes and that a higher MIC, at least in a subset of strains, correlates with a specific mutational signature in the gene *hsmA*.

## Materials and methods

### Strain characterisation

The strains analysed here were collected as part of the COMBACTE-CDI study. Strains were characterised by PCR ribotyping following the *C. difficile* Ribotyping Network of England and Northern Ireland protocol and tested for metronidazole susceptibility in agar dilution experiments at the Healthcare Associated Infection group of the University of Leeds, United Kingdom. ^16, 17^ Confirmation of PCR ribotype and additional susceptibility testing was performed at the Netherlands Reference Laboratory of *C. difficile*, hosted at the Leiden University Medical Center, The Netherlands. ^18^ Use of the strains for the present study was approved by the Management Board of the COMBACTE-CDI Consortium. Strains were further characterised by whole genome sequencing (see below).

Metronidazole minimal inhibitory concentrations were determined using the agar dilution method according to CLSI guidelines on either Wilkins-Chalgren agar (in Leeds) or Brucella Blood Agar (BBA) (in Leiden). ^19^ Although no formal breakpoints have been defined, we used the EUCAST epidemiological cutoff of 2 mg/L to define metronidazole resistance in *C. difficile* in this study. ^20^

### Determining medium-dependent metronidazole resistance

Agar plates on which *C. difficile* was grown were always incubated anaerobically at 37°C in a Don Whitley A55 workstation (10% CO_2_, 10% H_2_ and 80% N_2_ atmosphere). For determining medium-dependent metronidazole resistance, *C. difficile* strains were first grown on reduced Brain Heart Infusion (BHI) agar plates supplemented with 0,5% yeast extract (Sigma-Aldrich) and *Clostridium Difficile Selective Supplements* (CDSS, Oxoid) for 24-48 hours. From these plates bacterial suspensions corresponding to 2.0 McFarland turbidity were made in PBS. These suspensions were then applied on BBA plates (bioMérieux) or on BHI agar supplemented with 0.5% yeast extract and when applicable with 1 μg/ml vitamin K (Sigma-Aldrich), 5% sheep blood (Thermo Fisher Scientific) and/or 5 μg/ml hemin (Sigma-Aldrich). Metronidazole or vancomycin E-tests (bioMérieux) were then applied and growth was evaluated after 48 hours of anaerobic incubation. Pictures were taken with an Interscience Scan 500 Automatic colony counter.

### Whole genome sequencing and signature analysis

As part of the COMBACTE-CDI project, DNA extracts of the selected isolates were processed for whole genome sequencing at GSK Bio, as per their standard operating procedures. Briefly, total genomic DNA’s were quantitated using Quant-iT dsDNA High-Sensitivity Assay Kit (Life technologies) and SYNERGY H1 microplate reader. Sequencing libraries were prepared from 1 ng of DNA using the Nextera XT DNA Library Prep kit (Illumina) and libraries concentrations were normalised using bead normalisation as described by the manufacturer. Sequencing was performed on the Illumina MiSeq platform with MiSeq v3 600-cycles kit or on the Illumina NextSeq 500 platform with NextSeq 550 High-Output v 2.5 300-cycles kit. FASTQ files passed the quality control checks and strain identification confirmation (C. *difficile*), as performed in the BIOMERIEUX EPISEQ® CS beta platform (https://www.biomerieux-episeq.com/).

For the SNP analysis, reads were mapped to the *hsmA*, *hsmR*, *hatT* and *hatR* genes obtained from the reference sequence of strain R20291 (GenBank entry FN545816.1). Mapping of reads to the reference sequences was done with Bowtie2 tool (version 2.3.4, options: --local --qc-filter) and results of this mapping were analysed using Samtools (version 1.7, default options) and depth of the reads coverage was calculated using IGV tool (version 2.3.98, options: -w 1). ^21–23^ Average depth calculated for all positions of the analysed isolates is shown in Supplementary Table 1. Values of depth of coverage obtained for each position (nucleotides or indels) for all samples were filtered using a minimum coverage of 5 reads for each position. Mapping results for all positions were compared between all strains. Positions where at least one strain has a mutation or variation compared to the corresponding reference sequence were included in the generation of the genetic signature of *hsmA, hsmR, hatT* and *hatR*.

### Phylogenetic analysis

ST15 and 15-like publicly available whole-genome sequencing data (Supplemental Table 2) were selected based on a previous analysis, and corresponding FASTQ files were downloaded from the SRA (n=85). ^24^ wgSNP analysis was performed on the Enterobase platform with a selected minimum frequency threshold of 0.1. ^25, 26^ Three entries were excluded from the analysis, as they did not pass the quality control of the Enterobase platform (Supplemental Table 2). ^26^ A wgSNP maximum likelihood tree was generated with RaxML and tree branches were represented in a log scale for clarity. ^27^

## Results

### Metronidazole resistance is observed in the COMBACTE-CDI strain collection

The Combatting Bacterial Resistance in Europe – *Clostridioides difficile* infections (COMBACTE-CDI) is a multi-centre European-wide project with an aim to provide detailed understanding of the epidemiology and clinical impact of CDI across the whole healthcare economy in Europe. Sites testing both in-patient and community samples were recruited from 12 countries across Europe. All diarrheal faecal samples (regardless of tests requested by physician) were submitted to a central laboratory (Leeds, UK) on two selected days between July and November 2018. From these samples *C. difficile* was isolated and tested by PCR ribotyping. The metronidazole Minimum Inhibitory Concentration (MIC) for 213 clinical isolates (Belgium n=3, France n=4, Greece n=4, Ireland n=1, Italy n=23, Netherlands n=8, Poland n=29, Romania n=37, Slovakia n=1, Spain n=43, Sweden n=12, UK n=48) were determined by Wilkins-Chalgren agar dilution. ^17^ Of these, 22 isolates (10%) were found to be resistant to metronidazole using the EUCAST criteria as a cut-off (MIC≥2 mg/L) and were sent to the *C. difficile* reference laboratory of the Leiden University Medical Center for further study. ^20^ When possible, an RT-matched isolate with an MIC<2 mg/L from the COMBACTE-CDI collection was also provided. PCR ribotypes submitted were RT002 (n=3); RT010 (n=7); RT016 (n=1); RT018 (n=3); RT027 (n=12); RT176 (n=1); RT181 (n=4) and RT198 (n=1) (Table 1).

**Table 1.** Characteristics of the strains from this study. MIC refers to the minimal inhibitory concentration for metronidazole as determined by agar dilution at the Leiden University Medical Center (see Materials and Methods).

| Isolate | Location of participant at time of sample | Country of origin | PCR ribotype | MIC (mg/L) |
| --- | --- | --- | --- | --- |
| GSK234 | Inpatient | Sweden | 002 | 0.25 |
| GSK6 | Community doctor | UK | 002 | 0.25 |
| GSK7 | Inpatient | UK | 002 | 0.125 |
| GSK22bis | Inpatient | UK | 010 | 0.125 |
| GSK211 | Inpatient | Romania | 010 | 0.25 |
| GSK241 | Inpatient | France | 010 | 4 |
| GSK242new | Inpatient | France | 010 | 4 |
| GSK246 | Inpatient | France | 010 | 0.5 |
| GSK313 | Inpatient | Spain | 010 | 4 |
| GSK184 | Inpatient | Poland | 016 | 2 |
| GSK39 | Community doctor | Italy | 018 | 0.5 |
| GSK302 | Inpatient | Italy | 018 | 0.5 |
| GSK303 | Inpatient | Italy | 018 | 0.25 |
| GSK54 | Inpatient | UK | 027 | 0.25 |
| GSK55 | Inpatient | Romania | 027 | 0.25 |
| GSK60 | Inpatient | Poland | 027 | 0.5 |
| GSK61 | Inpatient | Poland | 027 | 0.5 |
| GSK62 | Inpatient | Poland | 027 | 0.5 |
| GSK63 | Inpatient | Poland | 027 | 0.5 |
| GSK64 | Inpatient | Poland | 027 | 0.5 |
| GSK65 | Inpatient | Poland | 027 | 0.5 |
| GSK179 | Inpatient | Poland | 027 | 0.5 |
| GSK318 | Inpatient | Poland | 027 | 0.5 |
| GSK325 | Inpatient | Poland | 027 | 0.5 |
| GSK327 | Inpatient | Poland | 027 | 0.5 |
| GSK258 | Inpatient | Slovakia | 176 | 0.5 |
| GSK110 | Inpatient | Romania | 181 | 0.5 |
| GSK113 | Inpatient | Romania | 181 | 0.25 |
| GSK114 | Inpatient | Poland | 198 | 0.5 |
| GSK180 | Community doctor | Romania | 181 | 0.5 |
| GSK190 | Inpatient | Romania | 181 | 0.25 |

As metronidazole is generally quite rare, our findings underscore the importance for investigating large collections of clinical isolates to enrich for strains that are resistant, in order to investigate possible underlying causes of resistance.

### Low level resistance to metronidazole is not due to carriage of the pCD-METRO plasmid

We further investigated the clinical isolates from the COMBACTE-CDI strain collection (n=32; Table 1). To correct for interlaboratory differences and to make the results directly comparable to our previous study, ^10^ we performed PCR ribotyping and antimicrobial susceptibility testing by agar dilution according to CLSI standards on Brucella Blood agar (BBA) in a second laboratory. ^10, 28^ A single strain showed discrepant results in PCR ribotyping and was therefore excluded from further analysis. We found 4/31 strains resistant to metronidazole (13% of the preselected isolates, Table 1). Three of these 4 strains belonged to RT010 (MIC=4 mg/L) and the fourth isolate belonged to RT016 (MIC=2 mg/L). All of these strains were also identified as resistant in the initial susceptibility testing in Leeds. The interlaboratory difference in antimicrobial susceptibility may be explained by differences in testing methodology, but were not further investigated here.

To determine whether the observed resistance was due to the presence of the pCD-METRO plasmid, we performed a reference assembly of the sequence reads obtained from whole genome sequencing of these isolates against the pCD-METRO reference sequence (obtained from the European Nucleotide Archive BioProject number PRJEB24167). No reliable mapping of reads to the reference sequence was found and – in line with this finding – the strains were negative in a PCR assay directed against pCD-METRO (data not shown). ^10^ We therefore conclude that these strains do not carry pCD-METRO and that a different mechanism confers metronidazole resistance in these isolates.

### Resistance to metronidazole is medium dependent

We have previously described a strain that demonstrated medium-dependent metronidazole resistance (MIC=4 mg/L) independent of pCD-METRO. ^10^ In order to test if the metronidazole resistant phenotype of the four strains from the COMBACTE-CDI study could similarly be medium-dependent, strain GSK241 (MIC=4 mg/L) was plated on BHI agar that is routinely used in our laboratory, BBA (containing 5% defibrinated sheep blood, 1 μg/ml vitamin K and 5 μg/ml hemin), and BHI blood agar (BHI agar supplemented with 5% defibrinated sheep blood, 1 μg/ml vitamin K and 5 μg/ml hemin) after which a metronidazole E-test was applied. We found that strain GSK241 was susceptible to metronidazole (MIC=0.25 mg/L) on BHI medium but resistant (MIC≥2 mg/L) on both BBA agar and BHI blood agar (Figure 1). These results indicate that components present in blood agar are responsible for this medium-dependent resistant phenotype.

**Figure 1.**
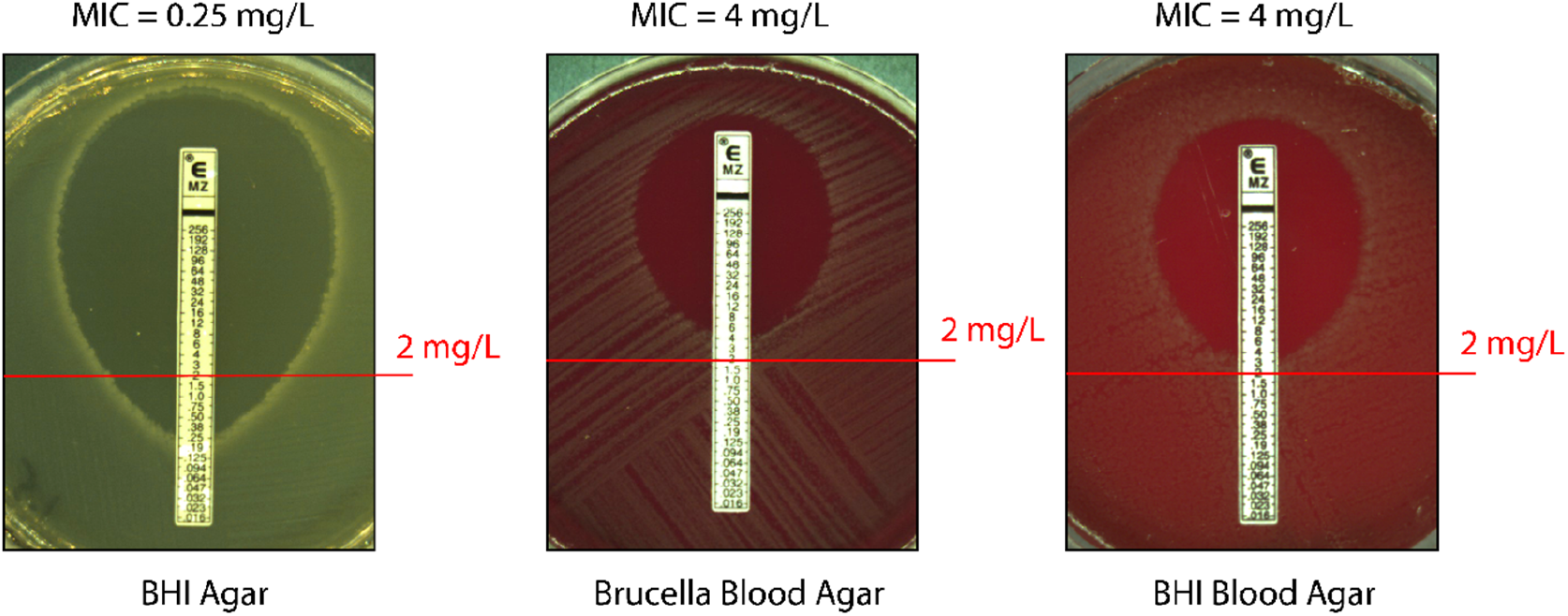
Medium-dependent resistance to metronidazole is independent of base broth. Strain GSK241 was grown and resuspended to 2.0 McFarland turbidity and spread on BHI agar, Brucella Blood Agar (supplemented with 1 μg/ml vitamin K and 5 μg/ml hemin) and BHI Blood Agar (BHI broth supplemented with 5% sheep blood, 1 μg/ml vitamin K and 5 μg/ml hemin). An E-test was placed and plates were incubated for 48h prior to imaging. Red lines indicate the epidemiological cut-off value for metronidazole as determined by EUCAST and have been used to determine resistance in this study ^20^. Numbers between brackets at the top of the image correspond to the reported MIC value on BHI agar, Brucella blood agar and BHI blood agar, respectively.

We wondered whether the medium-dependent change in MIC values was a general characteristic of *C. difficile* irrespective of the resistance phenotype, or specific to the resistant strains. For this reason, we tested selected COMBACTE-CDI strains by E-test on both BHI agar and BBA (Table 2). All strains were clearly susceptible on BHI agar (MIC < 0,5 mg/L) but, with the exception of GSK234, showed a 4-to-32-fold increase in MIC when tested on BBA compared to BHI agar. This medium-dependent increase in MIC was not restricted to a specific RT, as the phenotype was seen for strains belonging to diverse types (RT010, RT016, RT018, RT027, RT176, RT181 and RT198).

**Table 2.** Metronidazole minimal inhibitory concentrations on different agar media as determined by E-test. Strains were tested on BHI agar, BHI agar supplemented with 5 μg/ml hemin or BBA as determined by E-test. ND = not determined. MIC = minimal inhibitory concentration of metronidazole under the conditions tested

| Strain name | PCR Ribotype | E-test MIC BHI | E-test MIC BBA | E-test MIC BHI + hemin |
| --- | --- | --- | --- | --- |
| GSK234 | 002 | 0.064 | 0.064 | 0.064 |
| GSK241 | 010 | 0.25 | 4 | 6 |
| GSK242new | 010 | 0.25 | 6 | 6 |
| GSK313 | 010 | 0.38 | 4 | 8 |
| GSK246 | 010 | 0.125 | 1 | 1 |
| GSK184 | 016 | 0.25 | 2 | ND |
| GSK39 | 018 | 0.125 | 1.5 | 1 |
| GSK318 | 027 | 0.125 | 1 | 1 |
| GSK327 | 027 | 0.125 | 2 | 2 |
| GSK325 | 027 | 0.125 | 1.5 | ND |
| GSK60 | 027 | 0.125 | 4 | ND |
| GSK61 | 027 | 0.125 | 2 | 1.5 |
| GSK62 | 027 | 0.094 | 1.5 | ND |
| GSK63 | 027 | 0.125 | 1.5 | ND |
| GSK64 | 027 | 0.125 | 1.5 | 1.5 |
| GSK65 | 027 | 0.094 | 1 | ND |
| GSK179 | 027 | 0.125 | 1.5 | ND |
| GSK258 | 176 | 0.16 | 1 | 1.5 |
| GSK110 | 181 | 0.125 | 1 | 1.5 |
| GSK180 | 181 | 0.094 | 1 | 1 |
| GSK114 | 198 | 0.125 | 2 | 2 |

Taken together, our data suggest that components present in BBA/BHI blood agar result in reduced susceptibility of strains to metronidazole through a general mechanism. In the case of strains GSK184, GSK241, GSK242NEW and GSK313 this leads to these strains being qualified as metronidazole resistant.

### Heme is required for medium-dependent differences in metronidazole susceptibility

The fact that the medium-dependent increase in MIC was observed for both BBA and BHI blood agar (Figure 1) suggests that the phenotype is independent of the base broth and is likely to be mediated by the supplementation with vitamin K, hemin, and/or blood.

For practical purposes, we evaluated the effect of hemin on the metronidazole resistance phenotype of strains GSK64 (metronidazole susceptible) and GSK241 (metronidazole resistant). We found that supplementation of BHI with 5 μg/ml hemin raised the metronidazole MIC to levels similar to those observed for the BBA and BHI blood agar plates (Table 2; Figure 2A).

**Figure 2.**
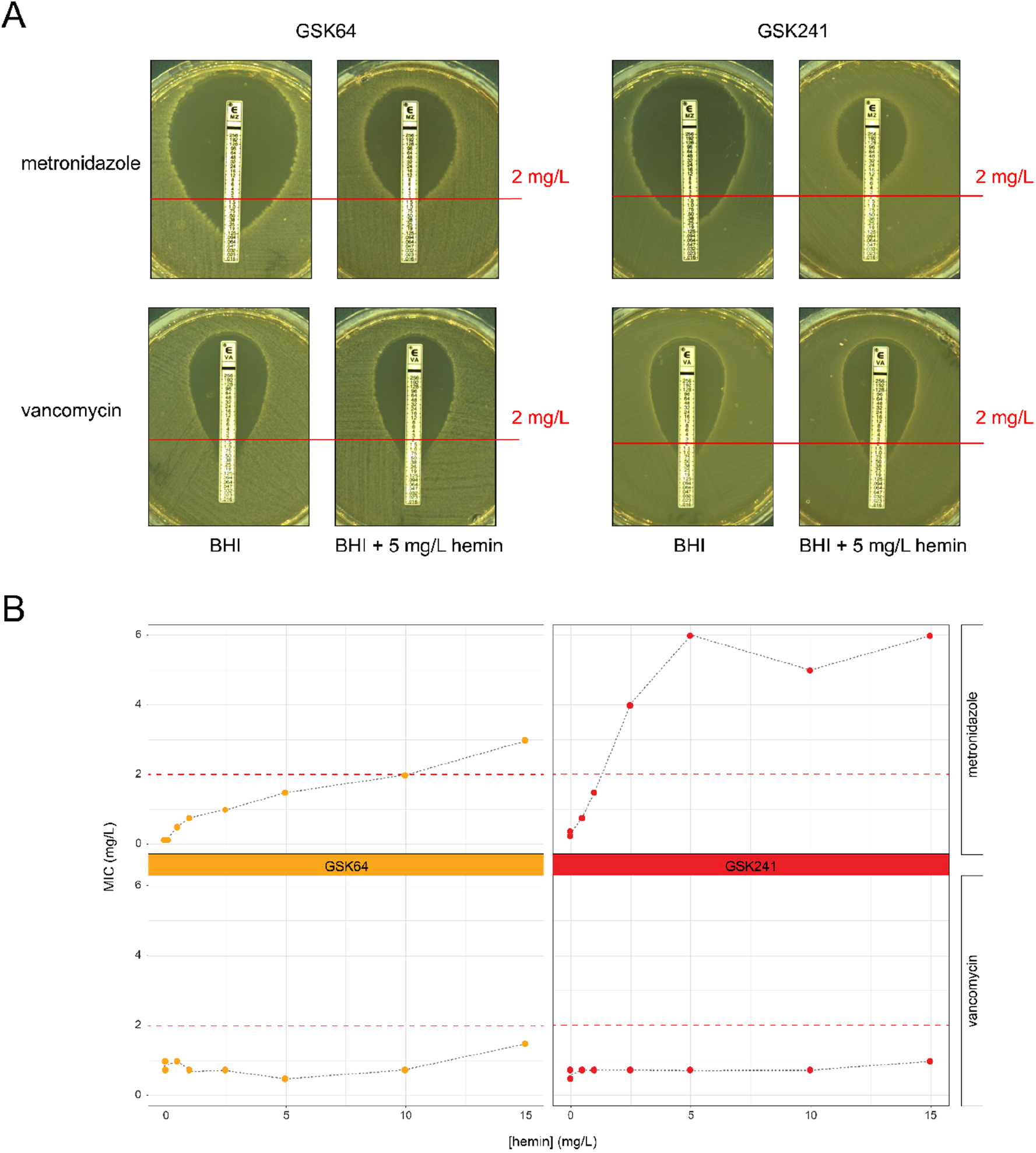
Heme supplementation increases the MICs to metronidazole, but not to vancomycin. **A**) Strains GSK64 and GSK241 were resuspended in PBS to 2.0 McFarland turbidity and plated on BHI agar and on BHI agar supplemented with 5 μg/ml hemin. Subsequently E-tests for metronidazole and vancomycin were applied and plates were incubated for 48h prior to imaging. Red lines indicate the epidemiological cut-off value for metronidazole as determined by EUCAST and have been used to determine resistance in this study ^20^. For a complete overview of the MICs to metronidazole of these strains on BHI agar and BHI agar supplemented with hemin, see Table 2. **B**) Typical metronidazole (top) and vancomycin (bottom) E-test results performed on strains GSK64 (orange) and GSK241 (red) and when grown on BHI supplemented with various concentration of hemin (0; 0.5; 1; 2.5; 5; 10; 15 mg/L). The dashed line indicates the epidemiological cut-off value for metronidazole and vancomycin as determined by EUCAST (2 mg/L).

We extended this finding to a selection of COMBACTE-CDI strains and found that, with the exception of strain GSK234, all tested strains showed an 8-to-24-fold increase in metronidazole MIC on BHI supplemented with hemin compared to BHI alone (Table 2). These results mirror those obtained for the E-test on BBA, indicating that hemin is the main determinant of medium-dependent differences in metronidazole MIC for these strains. Under our experimental conditions, the hemin-dependent increase in MIC appears to be specific to metronidazole, as no increase in MIC was observed for vancomycin (Figure 2A).

We next assessed the MIC of both metronidazole and vancomycin of strains GSK64 and GSK241 with a range of 0-15 μg/ml hemin. Both strains show a gradual increase in MIC for metronidazole that in the case of GSK241 saturates at > 5μg/ml hemin; the MIC for GSK64 appears to increase further at higher concentration of hemin. In contrast, no increase in vancomycin MIC was seen even in the presence of the highest concentrations of hemin tested (Figure 2B).

Altogether these results demonstrate that the presence of heme is crucial for a medium-dependent resistance phenotype in *C. difficile* and that this appears to be specific for metronidazole.

### An *hsmA* genetic signature is associated with increase metronidazole MICs in PCR ribotype 010 strains

Recent work has shown that four genes (*hsmA*, *hsmR*, *hatT*, *hatR*) are differentially regulated in response to heme, and that the products of the *hsmAR* operon improve growth of a *C. difficile* PCR ribotype 027 strain in the presence of metronidazole. ^29, 30^ For this reason we performed a single nucleotide polymorphism (SNP) analysis on the genes *hatR*, *hatT*, *hsmR* and *hsmA* using the sequences from the RT027 strain R20291 (GenBank accession FN545816) as a reference.

Using variant positions, we identified signatures for these genes for each strain from the COMBACTE-CDI collection. In the case of *hatR*, *hatT* and *hsmR* these signatures were conserved within a PCR ribotype, and even across closely related PCR ribotypes (e.g RT027, RT198, RT181 and RT176 share the same signatures) (Supplementary Table 1). However, for *hsmA,* we observed two distinct but related signatures within a single PCR ribotype. SNPs were identified in the 399-bp *hsmA* gene at positions 129, 249, 366, 372 and 392, resulting in a 5-base pair signature sequence. This signature is GGCAT for the RT027 and RT027-like isolates, TGTAC in RT002 and RT018 isolates and TATAC in some RT010 isolates (Figure 3A). Interestingly, the signature sequence TAT-C was found in all 3 metronidazole resistant RT010 isolates. The deletion at position 372 results in a frameshift and alters the primary amino acid sequence of the C-terminus of the HsmA protein.

**Figure 3.**
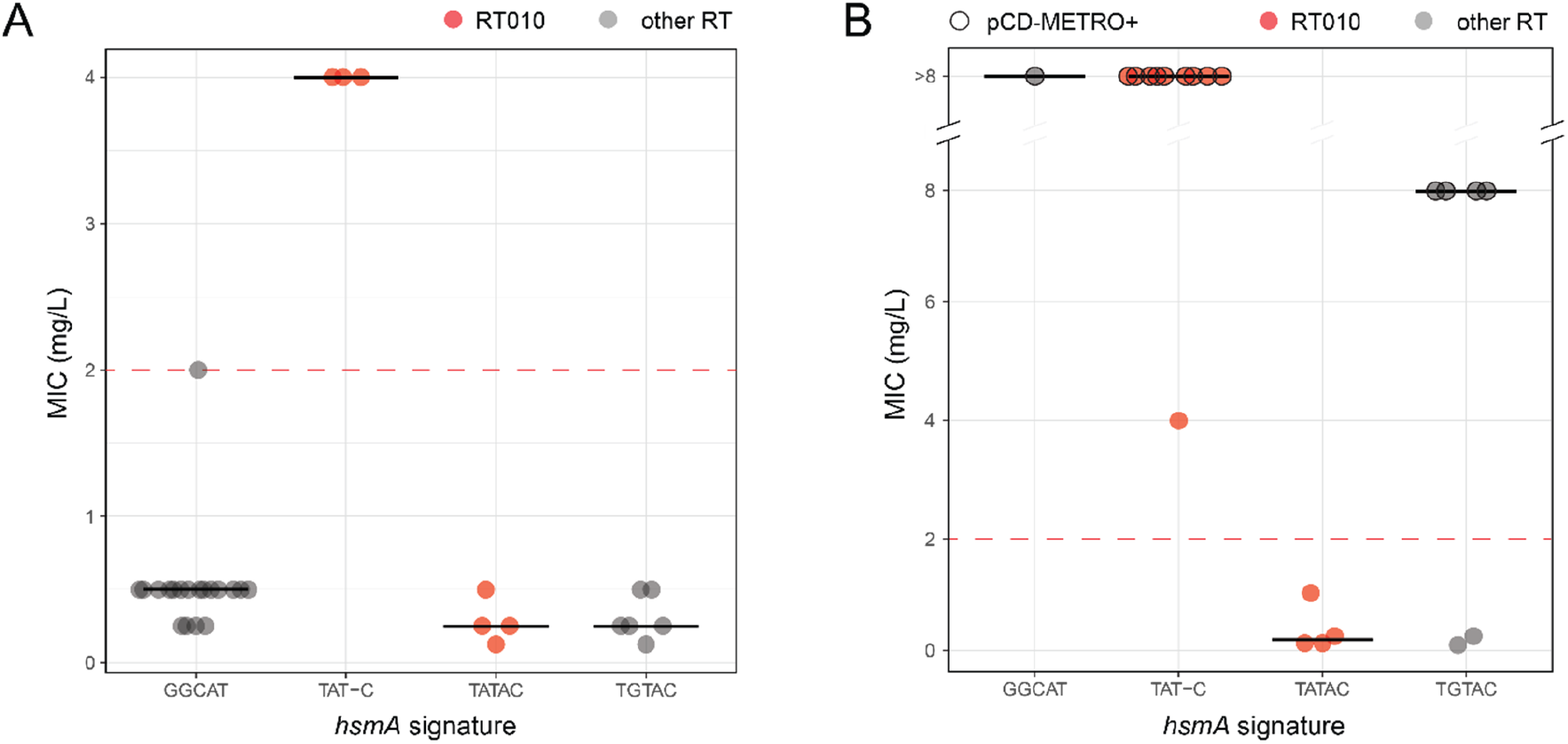
TAT-C signature in *hsmA* correlates to metronidazole resistance in RT010 isolates. The horizontal black bars represent the median MIC. **A**) *hsmA* signature sequences based on SNP analysis in the COMBACTE-CDI clinical isolates. Strains containing the GGCAT signatures belong to RT016, RT027, RT176, RT181 and RT198. The TAT-C and TATAC signatures correspond to RT010 in this study, whereas RT002 and RT018 containing sequence TGTAC. **B**) *hsmA* signature based on SNP analysis in the clinical isolates sequenced in the pCD-METRO study ^10^. The signature sequence GGCAT is found in RT027, TAT-C and TATAC in RT010 and TGTAC in RT012 and RT020.

In order to validate the significance of the signature, we performed the same *hsmA* SNP analysis on whole genome sequences obtained from another collection of isolates enriched for metronidazole resistance. ^10^ This collection contains RT010, RT020 and RT027 isolates which have been characterised with respect to metronidazole MIC and pCD-METRO carriage. We expected to find a similar clustering of *hsmA* signature sequence and PCR ribotype and predicted that the previously described RT010 isolate showing medium-dependent resistance (MIC=4 mg/L) would carry the 1-bp deletion in *hsmA*. Indeed, we found this was the case (Figure 3B). Strikingly, all highly resistant RT010 strains that carried pCD-METRO also contained the TAT-C *hsmA* signature sequence (Figure 3B).

We wondered how frequently this deletion could be found in RT010 isolates, as metronidazole resistance is most commonly observed in this RT. For this reason, we performed whole genome SNP (wgSNP) analysis through the Enterobase platform on sequences available in the Sequence Read Archive (SRA) of multilocus sequence type 15 (ST15; which includes RT010) (n=57) and ST15-like strains (n=4) as well as the sequences of the RT010 isolates described earlier in this study (n=21) ^10,^ 24-26. We found that the TAT-C signature in *hsmA* was detected in a specific lineage of ST15- and ST15-like strains originating from different countries (Figure 4). One out of 61 (1.6%) of the ST15- and ST15-like isolates was found to contain the 1-bp *hsmA* deletion (accession number ERR125985), but no metadata was available for this strain in the SRA to confirm metronidazole resistance. Interestingly, pCD-METRO carriage is distributed throughout the lineage with the TAT-C *hsmA* signature, suggesting that pCD-METRO may be preferentially acquired in strains with pre-existing low-level metronidazole resistance.

**Figure 4.**
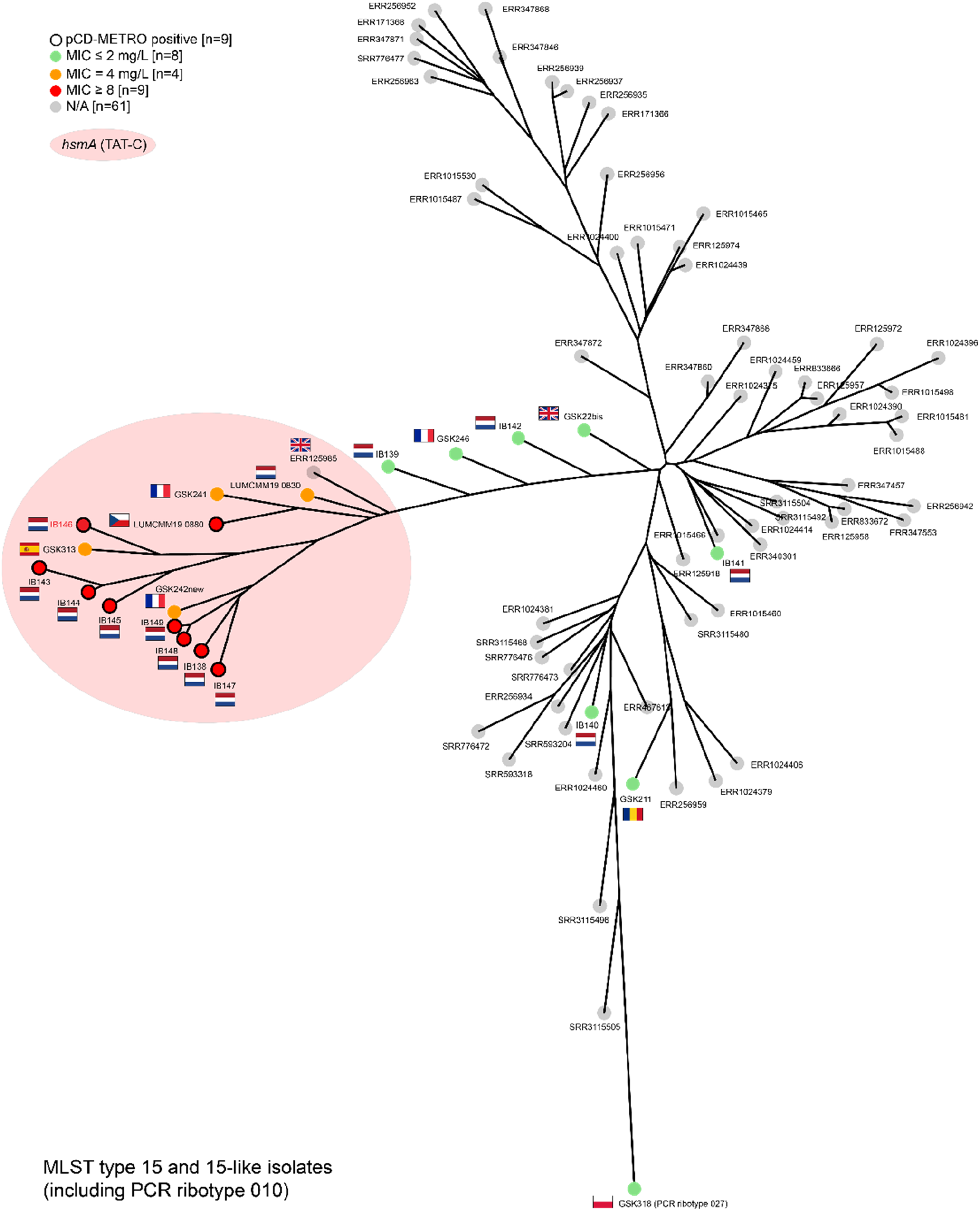
The C-terminal deletion in *hsmA* is associated with a lineage of ST15 and ST15-like isolates. When available, metronidazole susceptibility data and pCD-METRO carriage are indicated as illustrated in the legend. Country flags prior to strain name indicate the strain origin, when known. Strain names^10^ or SRA accession numbers are included. Distances in tree are shown in logarithmic scale.

Altogether our results demonstrate that a specific signature of *hsmA*, resulting in an altered C-terminal protein sequence, is associated with heme-dependent metronidazole resistance as well as pCD-METRO carriage in RT010 strains of *C. difficile*.

## Discussion

In this study we describe a collection of clinical *C. difficile* isolates that demonstrate a heme-dependent increase in MIC to metronidazole and make the observation that four strains determined to be resistant to metronidazole required heme supplementation for this phenotype. Additionally, we show that a C-terminal deletion at position 372 in *hsmA* correlates to metronidazole resistance in RT010 isolates.

The observation that the MICs for certain antibiotics vary depending on the type of medium used has been well documented in other organisms, but little to no data is available for *C. difficile*. ^*10, 31*^ One of the best known examples is the effect of divalent calcium on daptomycin susceptibility of a variety of organisms but other examples have been documented as well. ^32^ For instance, *Escherichia coli* was sensitive to bleomycin in LB broth, but was resistant to this antibiotic in glucose minimal medium, though the mechanism behind this difference remains unclear. ^33^ Similar results were obtained for *Moellerella wisconsensis* and *Proteus* spp for fosfomycin resistance when comparing MICs on IsoSensitest broth and (cation-adjusted) Mueller Hinton broth. ^34, 35^ Additionally, medium-dependent activity of gentamicin sulfate against enterococci has also been encountered, showing medium-dependent activity of antibiotics against bacteria is not a phenomenon restricted to gram-negative organisms. ^36^

In this study we observed that almost all strains showed an increase in metronidazole MIC when grown on blood agar compared to BHI (Figure 1, Table 2). We note the limitation that the COMBACTE-CDI collection we analysed here does not encompass an unbiased collection of PCR ribotypes, and in particular clade 2 strains (RT016, RT027, RT176, RT181, RT198) appear to be overrepresented. We found that heme supplemented to the blood agar plates was the causative determinant for this phenotype (Figure 2, Table 2), presumably through the ability of heme to detoxify the nitro-radicals generated by metronidazole activation. ^37^ As lethal concentrations of antimicrobials are thought to generate toxic radicals by altering cellular metabolism, we expected to find heme-dependent alterations in antibiotic susceptibility for other antibiotics than metronidazole. 38, 39 However, no heme-dependent reduction in vancomycin susceptibility was found in our study, suggesting that the effect of heme shows specificity for metronidazole under the conditions tested.

Elegant work by Knippel *et al*. has demonstrated that reduced metronidazole susceptibility upon heme supplementation in R20291 was largely mediated by the *hsmRA* operon, leading to the question whether presence/absence or sequence variants of these genes underlie heme-dependent resistance in other RTs. ^29^ Based on the present dataset, however, we were unable to identify specific sequence variants of this operon (or in the *hatRT* operon also involved in heme detoxification) that could explain why the vast majority of strains are less susceptible to metronidazole upon heme supplementation. The genes appear to be (near-)universally conserved amongst different *C. difficile* types (data not shown) and the same signature is found in strains that do or do not respond to heme supplementation, and those that do or do not qualify as resistant (Table 1, Table 2, Supplemental Table 1). These results imply that other factors than the genome sequence of the *hatRT* and *hsmRA* operons can contribute to heme-dependent reduction in metronidazole susceptibility.

Nevertheless, we did identify a C-terminal deletion in *hsmA* that correlated to heme-dependent metronidazole resistance in RT010 (ST15) isolates (Figure 3, Figure 4). Isolates without this deletion did become less susceptible to metronidazole in presence of heme, but did not exceed the EUCAST criterium for resistance to this antibiotic. ^20^ We validated our findings in the collection of clinical isolates used in the pCD-METRO study. ^10^ At present, there is no structural information on the HsmA protein, though homology between the protein and heme-containing cytochromes has been noted. 29 HsmA has been postulated to act through sequestration of heme, but the effect of the altered C-terminal sequence on the affinity for heme remains to be elucidated.

The COMBACTE-CDI collection analysed here includes a limited number of strains per RT (Table 1). It will be interesting to see whether targeted analyses of larger collections of specific ribotypes will reveal additional sequence variants of *hsmAR* associated with reduced susceptibility or resistance to metronidazole.

Our data hint at a possible cumulative effect of chromosomal and extrachromomal determinants in metronidazole resistance as strains carrying the pCD-METRO plasmid are dispersed over the resistant ST15/ST15-like (RT010) lineage characterised by the TAT-C *hsmA* signature (Figure 4). Strains that possess both the C-terminal adenine deletion in *hsmA* and the pCD-METRO plasmid have a higher metronidazole MIC (MIC ≥ 8 mg/L vs 4 mg/L in the pCD-METRO negative RT010s). As no pCD-METRO positive RT010 isolate containing the TATAC signature sequence was present in this collection, we do not know if pCD-METRO carriage without the deletion can still result in an MIC of ≥ 8 mg/L, though this appears to be the case in RT020 and RT027. ^10^ Irrespective of the effect on metronidazole MICs, pCD-METRO carriage is associated with the 1-bp deletion in *hsmA* in RT010 isolates. Though this might result from a selection bias (by preferentially characterizing isolates with higher MICs), it is conceivable that the deletion facilitates pCD-METRO carriage in some way.

Our findings suggest that the heme-dependent reduction in metronidazole susceptibility is common in *C. difficile* (Table 2). Though heme levels can be elevated at the host-pathogen interface during CDI, ^30^ pathogenicity does not seem a requirement for the heme-dependent reduction in MIC as it is also found in non-toxigenic strains such as those belonging to RT010. For the same reason, it is unlikely that extensive and continued use of metronidazole provided the selective pressure for the acquisition and/or persistence of this phenomenon during evolution. ^6, 7, 40^ In the majority of the strains the MIC will likely not be raised over the EUCAST cut-off (2 mg/L) for resistance, ^20^ but they clearly do become less susceptible to metronidazole. Due to absorption in the small intestine and sequestration- or inactivation of the microbiota, levels of metronidazole at the end of the colon are potentially low as determined by concentrations found in fecal material. ^41–44^ It is therefore quite possible that a moderate increase in metronidazole MIC could in fact facilitate growth of *C. difficile* in patients treated with metronidazole. Whether or not heme-dependent reduced susceptibility plays a role in treatment failure (that does not appear to correlate with metronidazole resistance) remains to be established. ^10, 45^

An important implication of our findings is that a well-described testing method in diagnostic antimicrobial susceptibility testing is of the utmost importance. The type of media and supplementation used can influence the MIC of certain antibiotics as we have demonstrated for heme supplementation and metronidazole susceptibility. It is our experience that small interlaboratory differences in standard operating procedures exist despite conforming to CLSI standards, which may explain the differences in MICs sometimes found between institutions. As medium-dependent differences in antimicrobial susceptibility may be both antibiotic- and organism-specific, this argues for a standardised method per genus rather than a standard testing method for all anaerobic organisms.

In conclusion, we have demonstrated that heme is the causative agent of medium-dependent reduction in metronidazole susceptibility in clinical *C. difficile* isolates of different ribotypes, but does not influence vancomycin susceptibility. Additionally, we have found a deletion in the C-terminal part of *hsmA* which correlates to metronidazole resistance in RT010 isolates.

## Acknowledgements

We would thank the members of the COMBACTE-CDI Consortium for providing the *C. difficile* isolates and FASTQ files that were used in this study. The authors thank J. Corver and B. Hornung for helpful discussions.

## Transparency declaration

The COMBACTE-CDI consortium includes partners GSK, Pfizer, DaVolterra, AstraZeneca, Sanofi Pasteur and bioMérieux. The companies did not have a role in the design and execution of the experiments for this study but provided the WGS of the isolates (GSK) and allowed examination of the FASTQ files (bioMérieux). Conceived the study: IMB, EJK, JF, WKS. Performed experiments: IMB, CH, IMJGBS, BS, EC, JF. Analysed data: IMB, SN, IAS, VV, KD, JF, WKS. Drafted manuscript: IMB, WKS. All authors edited and approved the final version of the manuscript.

