## Supplemental Tables for "Heme is crucial for medium-dependent metronidazole resistance in clinical isolates of *C. difficile*"

| | | | *hsmA* | | *hsmR* | | *hatR* | | *hatT* | |
|---|---|---|---|---|---|---|---|---|---|---|
| Isolate | RT | MTZ MIC (mg/L) | Mean depth [SD] | Signature [129/249/366/372/392] | Mean depth [SD] | Signature [142] | Mean depth [SD] | Signature [45/132] | Mean depth [SD] | Signature [75/564/963/1035/1047/1173/1184/1665/1818/1914/1983/2028/2167/2219/2391/2421] |
| Reference (R20291) | | | | GGCAT | | G | | AA | | ACTGGCGACGCCTATC |
| GSK234 | 002 | 0.25 | 143.19 [43.21] | TGTAC | 95.84 [16.84] | A | 81.05 [30.74] | GG | 201.60 [66.52] | GTGGAAGATACCAGCT |
| GSK241 | 010 | 4 | 80.60 [20.48] | TAT-C | 58.29 [10.72] | A | 46.63 [11.74] | GG | 122.05 [36.68] | ATGAAATACACTTGCT |
| GSK242new | 010 | 4 | 65.26 [10.43] | TAT-C | 50.84 [21.74] | A | 45.26 [ 5.62] | GG | 106.10 [39.08] | ATGAAATACACTTGCT |
| GSK313 | 010 | 4 | 85.12 [20.43] | TAT-C | 49.11 [13.81] | A | 42.20 [10.61] | GG | 107.87 [40.15] | ATGAAATACACTTGCT |
| GSK246 | 010 | 0.5 | 65.97 [20.60] | TATAC | 65.36 [20.14] | A | 54.18 [ 9.11] | GG | 97.49 [27.34] | ATGAAATACACTTGCT |
| GSK184 | 016 | 2 | 51.83 [15.36] | GGCAT | 47.66 [10.70] | G | 38.37 [ 7.48] | AA | 89.39 [21.09] | ACTGGCGACGCCTATC |
| GSK39 | 018 | 0.5 | 91.28 [20.81] | TGTAC | 85.19 [22.99] | A | 94.80 [15.60] | GG | 130.78 [30.61] | ATGGAAGGTATCTGCT |
| GSK318 | 027 | 0.5 | 86.10 [15.79] | GGCAT | 63.28 [15.63] | G | 64.18 [ 7.77] | AA | 117.23 [35.19] | ACTGGCGACGCCTATC |
| GSK327 | 027 | 0.5 | 62.30 [12.78] | GGCAT | 64.69 [16.08] | G | 33.44 [ 4.27] | AA | 89.93 [26.66] | ACTGGCGACGCCTATC |
| GSK325 | 027 | 0.5 | 91.83 [21.36] | GGCAT | 73.75 [11.56] | G | 64.74 [ 7.91] | AA | 135.43 [43.32] | ACTGGCGACGCCTATC |
| GSK60 | 027 | 0.5 | 114.67 [22.67] | GGCAT | 65.90 [16.74] | G | 49.69 [13.68] | AA | 134.58 [44.03] | ACTGGCGACGCCTATC |
| GSK61 | 027 | 0.5 | 97.74 [30.19] | GGCAT | 48.62 [ 8.44] | G | 38.91 [ 7.13] | AA | 100.40 [41.67] | ACTGGCGACGCCTATC |
| GSK62 | 027 | 0.5 | 93.55 [28.49] | GGCAT | 48.37 [ 8.94] | G | 43.08 [10.86] | AA | 137.45 [46.61] | ACTGGCGACGCCTATC |
| GSK63 | 027 | 0.5 | 96.53 [16.07] | GGCAT | 82.08 [22.08] | G | 56.83 [ 5.03] | AA | 131.00 [41.78] | ACTGGCGACGCCTATC |
| GSK64 | 027 | 0.5 | 68.11 [ 8.71] | GGCAT | 66.69 [11.38] | G | 62.64 [ 6.53] | AA | 125.52 [33.71] | ACTGGCGACGCCTATC |
| GSK65 | 027 | 0.5 | 72.90 [16.61] | GGCAT | 57.54 [ 9.86] | G | 48.37 [ 9.09] | AA | 104.96 [31.48] | ACTGGCGACGCCTATC |
| GSK179 | 027 | 0.5 | 91.77 [23.85] | GGCAT | 58.82 [12.12] | G | 60.59 [ 6.56] | AA | 95.74 [21.97] | ACTGGCGACGCCTATC |
| GSK258 | 176 | 0.5 | 33.48 [ 7.77] | GGCAT | 21.03 [ 4.62] | G | 22.96 [ 2.93] | AA | 67.86 [23.06] | ACTGGCGACGCCTATC |
| GSK110 | 181 | 0.5 | 54.34 [14.05] | GGCAT | 33.00 [ 4.87] | G | 29.77 [ 4.86] | AA | 75.95 [19.59] | ACTGGCGACGCCTATC |
| GSK180 | 181 | 0.5 | 87.97 [27.13] | GGCAT | 69.74 [ 9.76] | G | 44.89 [15.60] | AA | 115.09 [28.72] | ACTGGCGACGCCTATC |
| GSK114 | 198 | 0.5 | 115.61 [38.37] | GGCAT | 83.66 [17.04] | G | 58.45 [11.90] | AA | 154.18 [49.23] | ACTGGCGACGCCTATC |

**Supplementary Table 1. NGS reads mapping to the reference sequences of the *hsmA*, *hsmR, hatT* and *hatT* genes and genetic signatures of the isolates.** MIC values indicate the results obtained from metronidazole agar dilution at the Leiden University Medical Center. RT = PCR ribotype. MTZ = metronidazole. MIC = minimal inhibitory concentration. SD = standard deviation of the mean. Bracketed numbers under Signature indicate variant base positions compared to the reference as described in Materials and Methods. Mismatches in the signatures with the corresponding positions in the reference sequences are shown with red shading.

| ST15 | | ST15-like |
|---|---|---|
| SRR776477 | ERR171368 | SRR3115496 |
| SRR776476 | ERR171366 | SRR3115480 |
| SRR776473 | ERR125974 | SRR3115468 |
| SRR776472 | ERR125972 | ERR340068 |
| SRR593318 | ERR125958 | ERR125985 |
| SRR593204 | ERR125957 | |
| SRR3115505 | ERR125918 | |
| SRR3115504 | ERR1024460 | |
| SRR3115492 | ERR1024459 | |
| ERR833672 | ERR1024439 | |
| ERR833666 | ERR1024414 | |
| ERR467613 | ERR1024406 | |
| ERR347872 | ERR1024400 | |
| ERR347871 | ERR1024396 | |
| ERR347868 | ERR1024390 | |
| ERR347866 | ERR1024381 | |
| ERR347860 | ERR1024379 | |
| ERR347846 | ERR1024375 | |
| ERR347553 | ERR1015530 | |
| ERR347457 | ERR1015501 | |
| ERR340301 | ERR1015498 | |
| ERR256963 | ERR1015490 | |
| ERR256959 | ERR1015488 | |
| ERR256956 | ERR1015487 | |
| ERR256952 | ERR1015481 | |
| ERR256942 | ERR1015471 | |
| ERR256939 | ERR1015466 | |
| ERR256937 | ERR1015465 | |
| ERR256935 | ERR1015460 | |
| ERR256934 | | |

**Supplemental Table 2**. ST15 and ST15-like isolates identified previously[1]. Entries marked in red did not pass the Enterobase[2] QC and were not included in Figure 4 of the main manuscript. The entry highlighted in green corresponds to an isolate that carries the TAT-C *hsmA* signature, but has not been characterized for metronidazole resistance.
